# Binding Behavior of Microbial Functional Amyloids on Solid Surfaces

**DOI:** 10.1101/2020.04.25.060962

**Authors:** Esra Yuca, Ebru Şahin Kehribar, Urartu Özgür Şafak Şeker

## Abstract

Self-assembling protein subunits hold great potential as biomaterials with improved functions. Among the self-assembled protein structures functional amyloids are promising unique properties such as resistance to harsh physical and chemical conditions their mechanical strength, and ease of functionalization. Curli proteins, which are functional amyloids of bacterial biofilms can be programmed as intelligent biomaterials. In order to obtain controllable curli based biomaterials for biomedical applications, and to understand role of each of the curli forming monomeric proteins (namely CsgA and CsgB from Escherichia coli) we characterized their binding kinetics to gold, hydroxyapatite, and silica surfaces. We demonstrated that CsgA, CsgB, and their equimolar mixture have different binding strengths for different surfaces. On hydroxyapatite and silica surfaces, CsgB is the crucial element that determines the final adhesiveness of the CsgA-CsgB mixture. On the gold surface, on the other hand, CsgA controls the behavior of the mixture. Those findings uncover the binding behavior of curli proteins CsgA and CsgB on different biomedically valuable surfaces to obtain a more precise control on their adhesion to a targeted surface.

## INTRODUCTION

Utilization of self-assembly abilities of protein subunits offers the fabrication of biomolecule-based new generation materials with improved functions. Collagen, actin, knot proteins, protein catenanes ^1^, viral capsid proteins ^2^, scaffold proteins ^3^, bacterial flagella ^4^, bacterial pili ^5^, and amyloids ^6^ are the examples of protein nanostructures that can self-assemble into an ordered supramolecular structure.

Amyloid proteins that are resistant to hard physical and chemical conditions can be classified into two groups as functional and pathological amyloids. Functional amyloids go beyond the misfolding and aggregation properties of pathologic amyloids and bring the useful properties of amyloids to the forefront. Pathologic amyloids are produced as a result of protein aggregation observed in neurodegenerative disease conditions while functional amyloids are produced to form biofilms by following a different developmental process than pathological amyloids. -

Biofilms are complex microorganism populations embedded in the extracellular matrix (ECM) that protect bacteria from adverse environmental conditions and formed by bacteria attaching to the surface. Resistance to external physical and chemical stresses increases in biofilms compared to planktonic life. Although the exact content of the biofilm matrix that encapsulates the bacteria varies depending on bacterial strains and environmental conditions, it is generally composed of proteins, nucleic acids, and polysaccharides.^7^ Bacterial biofilms, which are highly regulated, complex and dynamic systems, resistant to harsh environmental conditions with their ECM, have a great potential for the production and development of new generation materials.^8, 9^

In recent years there is an increasing trend towards the use of amyloids and amyloid-like fibrils as building blocks and as functional biomaterial systems ^9-14^. Functional amyloid curli fiber, which constitutes an important protein component of the biofilm extracellular biofilm matrix, is the most common bacterial amyloid protein genetically manipulated ^15, 16^. Transcription of bacterial curli amyloid nanofiber subunits is regulated by *csgBAC* and *csgDEFG* operons. Bacterial curli amyloid nanofibers consist of major CsgA and minor CsgB amyloid subunit proteins with N-terminal signal sequences. Ours and other’s studies have investigated the use of CsgA and CsgB amyloid subunits for bioinspired materials production and bioinspired processes or device development. In our previous study, we optimized the recombinant production and biochemical properties of CsgA and CsgB curli protein mutants ^17^. Controlling the production and secretion of CsgA and CsgB amyloid proteins is critical in terms of final mechanical characteristics of the biofilm-based material assembly. From this motivation, morphological and mechanical properties of CsgA and CsgB biofilm proteins have been investigated to provide stiffer or softer amyloid nanofiber assemblies ^18^. Bacterial biofilms were also used to form conductive protein nanofibers, integrating conductive peptide motifs for biofilm assembly in the previous work. ^18^ In another study, a series of genetic circuits were designed to control the secretion of CsgA-material binding peptide fusions upon extracellular signal processing of the available metallic nanoparticle precursors. ^19^. In addition to control the assembly of biofilm amyloids, patterning of them can also be controlled using genetic logic gates to build up organized living material system upon inducing the coexpression of different CsgA fibers^8^. Debenedictis et al. have proposed CsgA and CsgB structural models using protein structure prediction servers ^20^. Moser et al. have used light to pattern bacteria onto the materials such as textiles, ceramics, and plastic by controlling the expression of curli fibers.^21^ Besides, Zhang et al. have developed a coarse-grained curli fiber model to provide insight into adhesion mechanisms of curli subunit ^22^. In another study, HA coating has been prepared by a biofilm inspired approach using the CsgA component of *E. coli* biofilm and a short peptide bioinspired from dental biofilm^23^. Keten group has examined CsgA adsorption onto graphene and silica surfaces through atomistic simulation^24^. Since there is an increasing number of efforts to use biofilm proteins as intelligent materials ^18, 25-27^, there is a need for an extensive characterization of their interactions with solid surfaces. Here, we investigated binding kinetics of curli functional amyloids to biomedically important solid materials such as gold, hydroxyapatite, and silica, ^28-33^ In this study we aimed to understand the binding kinetic of the biofilm forming proteins on given solid surfaces to understand the strength of their binding in different settings. We observed that CsgB has a central importance to control the adhesiveness of the final CsgAB mix on silica and HA whereas the adhesiveness on gold surface Csga has a dominant effect. These clues can help one to control and limit biofilm formation on given surface by targeting genes encoding CsgB and CsgA proteins through genetic manipulation or adjusted gene copy numbers. This approach can be used to construct living biomaterials with tunable adhesive properties.

### Purification of CsgA and CsgB Curli Subunits

We cloned the region encoding the CsgA and CsgB proteins without the signal sequence in our previous study ^17^. CsgA and CsgB proteins were expressed and purified as previously described. Briefly, recombinant *E. coli* BL21 bacteria harboring CsgA and CsgB expressing plasmid constructs were grown in LB medium supplemented with ampicillin (100 mg/mL) at 37 °C. After overnight incubation, cell culture was diluted. At an OD600 of approximately 0.9, CsgA and CsgB expressions were induced with IPTG addition to a final concentration of 0.5 mM. Following two hours incubation, cells were harvested with centrifugation at 4000 g for 20 min, and cell pellets were suspended in phosphate buffer saline (PBS, 2.7 mM KCl, 137 mM NaCl, 2 mM KH2PO4, 10 mM Na2HPO4) at pH 7.0 that contained 6 M GdnHCl, 10 mM imidazole. Lysed cells were centrifuged at 10000 g for 20 min at 4 °C, and the remaining cellular debris was removed. A cobalt resin-based affinity method was applied for the purification of the proteins. The cobalt resin was first equilibrated with PBS lysis buffer. The supernatant was mixed with equilibrated resin, and unbound proteins were removed with centrifugation at 1000 g. After washing with the same buffer, CsgA and CsgB proteins were eluted from the resin with PBS buffer that contained 6 M GdnHCl, 150mM imidazole. CsgA and CsgB proteins were transferred to PBS buffer using centrifugal filter units (Thermo Scientific™ Pierce) with a molecular weight cutoff of 3 kDa. Purified protein oligomers were separated from monomers using 30 kDa centrifugal filter units (Thermo Scientific™ Pierce) for the following kinetic analyzes.

### Immunoblot Analysis

Purified CsgA and CsgB protein samples were treated with formic acid to break down protein oligomers and were freeze-dried to remove formic acid. Formic acid treated and lyophilized CsgA and CsgB proteins were dissolved in 2x SDS sample buffer. Protein concentrations were measured by the bicinchoninic acid (BCA) method. Final protein samples were analyzed by 15% sodium dodecyl sulfate-polyacrylamide gel electrophoresis (SDS–PAGE) followed by staining in Coomassie brilliant blue R-250 to detect the protein bands. Following SDS-PAGE under denaturing conditions, the proteins were transferred onto polyvinylidene difluoride (PVDF) membrane using a semi-dry electroblotting system (Bio-Rad). The blot was incubated in 5% non-fat dried milk in tris-buffered saline, 0.1% Tween 20 (TBS-T) at room temperature for an hour in order to block non-specific binding sites, then probed with 1:5,000 dilution of anti-his mouse primary antibody (Thermo Scientific Pierce) in TBS-T. After washing twice with TBS-T, the blot probed with 1:5,000 dilution of anti-mouse horseradish-peroxidase (HRP) conjugated secondary antibody (Thermo Scientific Pierce) in TBS-T. The membrane blot was rewashed with TBS-T and visualized with the Chemidoc MP imaging system (Bio-Rad) following an enhanced chemiluminescence (ECL) substrate (Bio-Rad) application.

### Quartz Crystal Microbalance (QCM-D)

CsgA, CsgB, and equimolar concentration mix of both proteins were incubated at room temperature for two days to induce fiber formation. All the protein samples were prepared in PBS buffer at concentrations of 1 µM, 2 µM, 2.5 µM, 3 µM, and 4 µM. After baseline equilibration using PBS buffer, CsgA, CsgB, and equimolar mix protein samples were introduced to QCM-D (QCM Q-Sense E1) chamber in sequential order. The absorption and material interaction behavior of the protein samples were evaluated on hydroxyapatite, silica, and gold surfaces in three experimental sets using a QCM-D system. After the delivery of each protein concentration to the system and the washing step, frequency and dissipation changes were recorded.

To characterize the adsorption of protein samples onto different sensor surfaces, data from three different overtone orders were analyzed using the Langmuir equilibrium model. This model relates the frequency change to the desorption strength by the equation Δ*f*= (*f*max x *C*)/(*k*d + *C*). *C* is the protein concentration, *f*maxvalue was estimated by the Langmuir isotherm, and *k*d value was determined by least-squares fitting.

Gibbs free energy is calculated using the formula ΔG°= -RT ln(k_eq_), where R is the gas equilibrium constant (1.987 kcal K ^-1^ mol ^-1^), T is the reaction temperature in K (298 K) and k_eq_ is 1/k_d_.

## Results and discussion

### Analysis of CsgA and CsgB monomers

Theoretical molecular weights of the recombinant CsgA and CsgB proteins were calculated as 15.52 kDa and 13.34 kDa, respectively, using the Swissprot Expasy tool. Purified CsgA and CsgB protein bands showed approximately at the same value compared to theoretical molecular weights in the Coomassie brilliant stained SDS-PAGE. Protein band were also verified using Western blotting (Figure S1).

### Kinetic Analysis

Curli, the adhesive protein that provides attaching to surfaces during biofilm formation, is an essential precursor for development of innovative materials with its adherent feature. We monitored the binding affinity of the protein polymers of CsgA, CsgB and their mixture for biomedically relevant surfaces namely gold, silica hydroxyapatite as presented in Figure 1.

**Figure 1.**
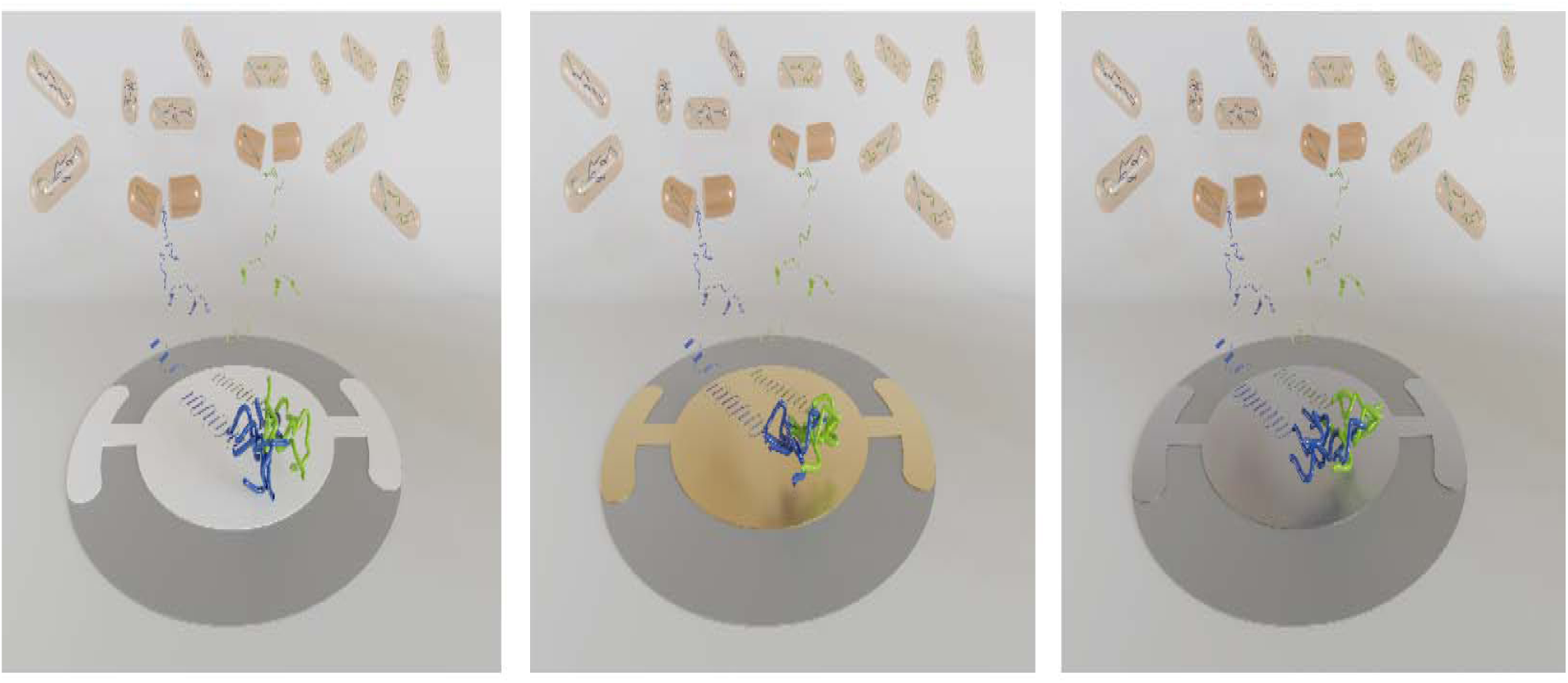
The interaction of CsgA polymer, CsgB polymer and their mixture with hydroxyapatite, gold and silica surfaces.

QCM-D is a powerful tool to track interactions of proteins with solid surfaces by providing real-time data for protein binding as well as dissipated energy upon protein / surface interaction. ^34, 35^.

The binding equilibrium constant and binding free energy of the proteins were calculated by measuring the frequency changes during protein addition processes and washing steps. Each of the polymerized CsgA, CsgB, and CsgAB proteins were deposited on three different types of QCM crystals, and the frequency change versus concentration graphs was drawn.

Upon addition of increasing concentrations of polymerized CsgA through the QCM-D system a drop in frequency for all three surfaces (Figure 2) was recorded. All incubated protein samples have highest binding affinity to HA surface compared to gold and silica. CsgB and CsgA-CsgB equimolar mixture polymers have higher binding affinity to gold than silica (Figure 3, Figure 4), while CsgA polymer exhibits approximately similar affinity for the two surfaces (Figure 2). Binding free energy of CsgA, CsgB, and CsgA-CsgB mix on gold, silica, and HA surfaces were calculated. (Figure 5).

**Figure 2:**
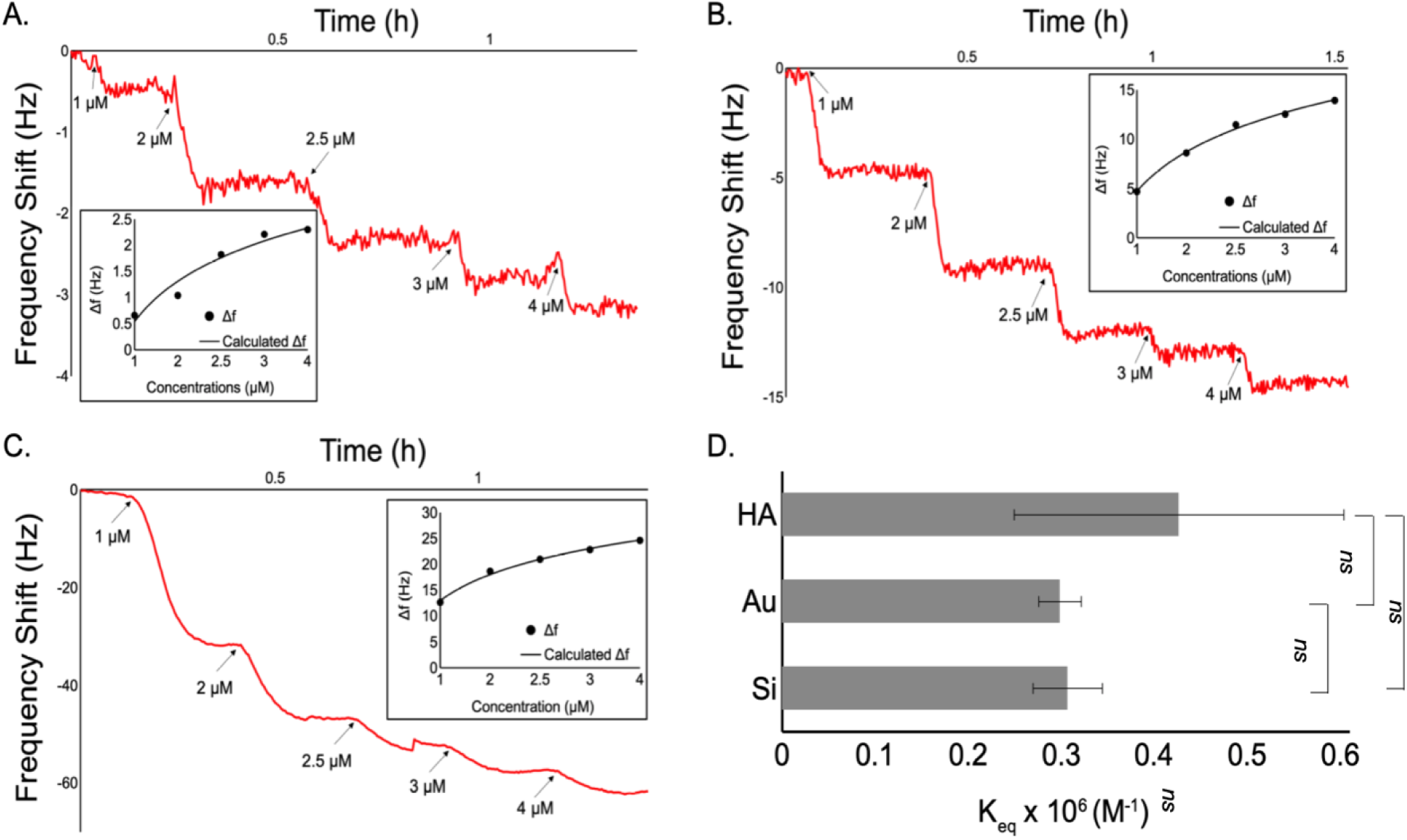
QCM measurement of aged CsgA. Binding kinetics to the (A) gold, (B) sili and (C) HA coated quartz surface is shown as a resonance frequency change. D. Binding equilibrium constants of polymerized CsgA on gold, silica, and HA coated surfaces.

**Figure 3.**
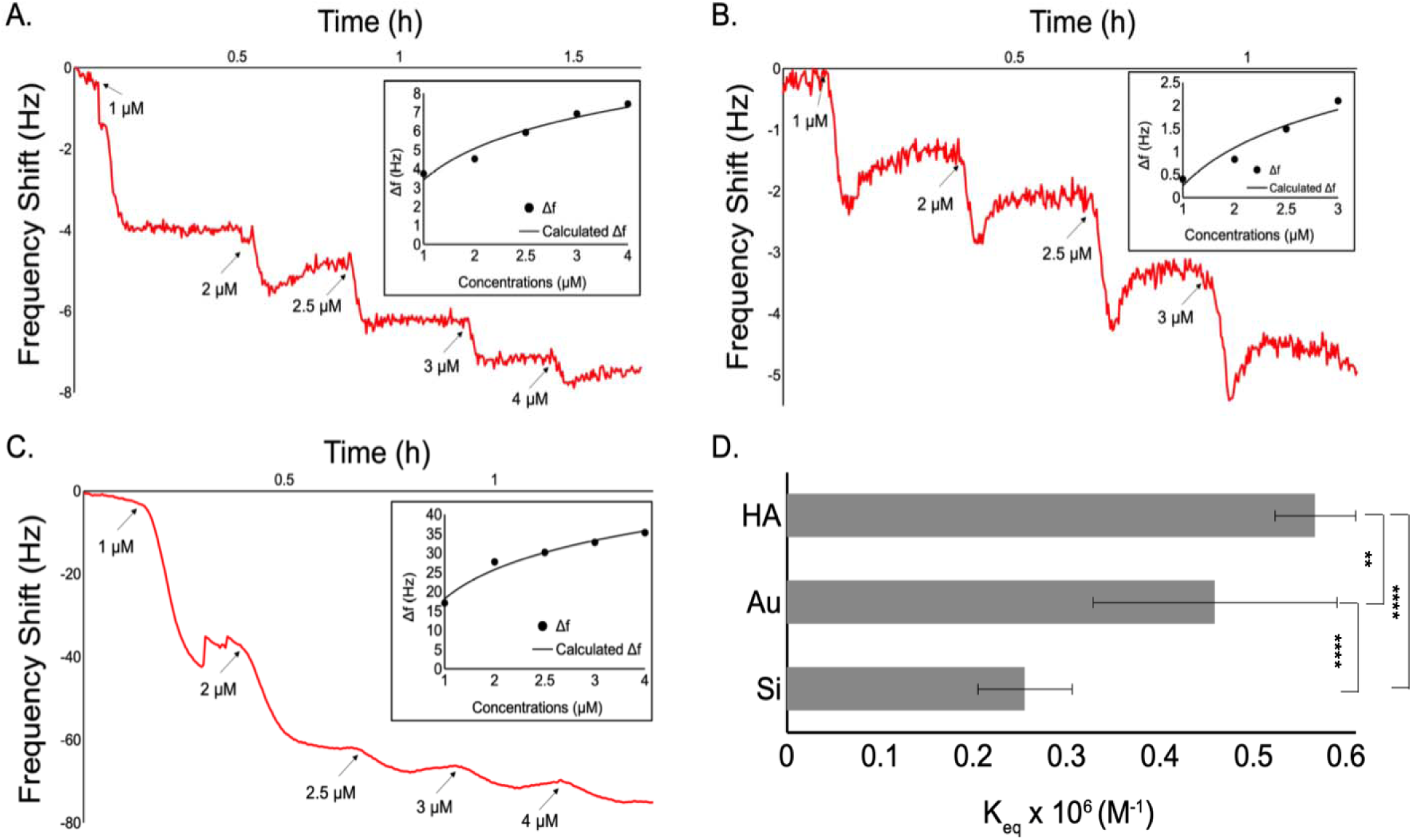
QCM measurement of incubated CsgB. Binding kinetics to the (A) gold, (B) silica, and (C) HA coated quartz surface is shown as a resonance frequency change. D. Binding equilibrium constants of polymerized CsgB on gold, silica, and HA coated surfaces.

**Figure 4.**
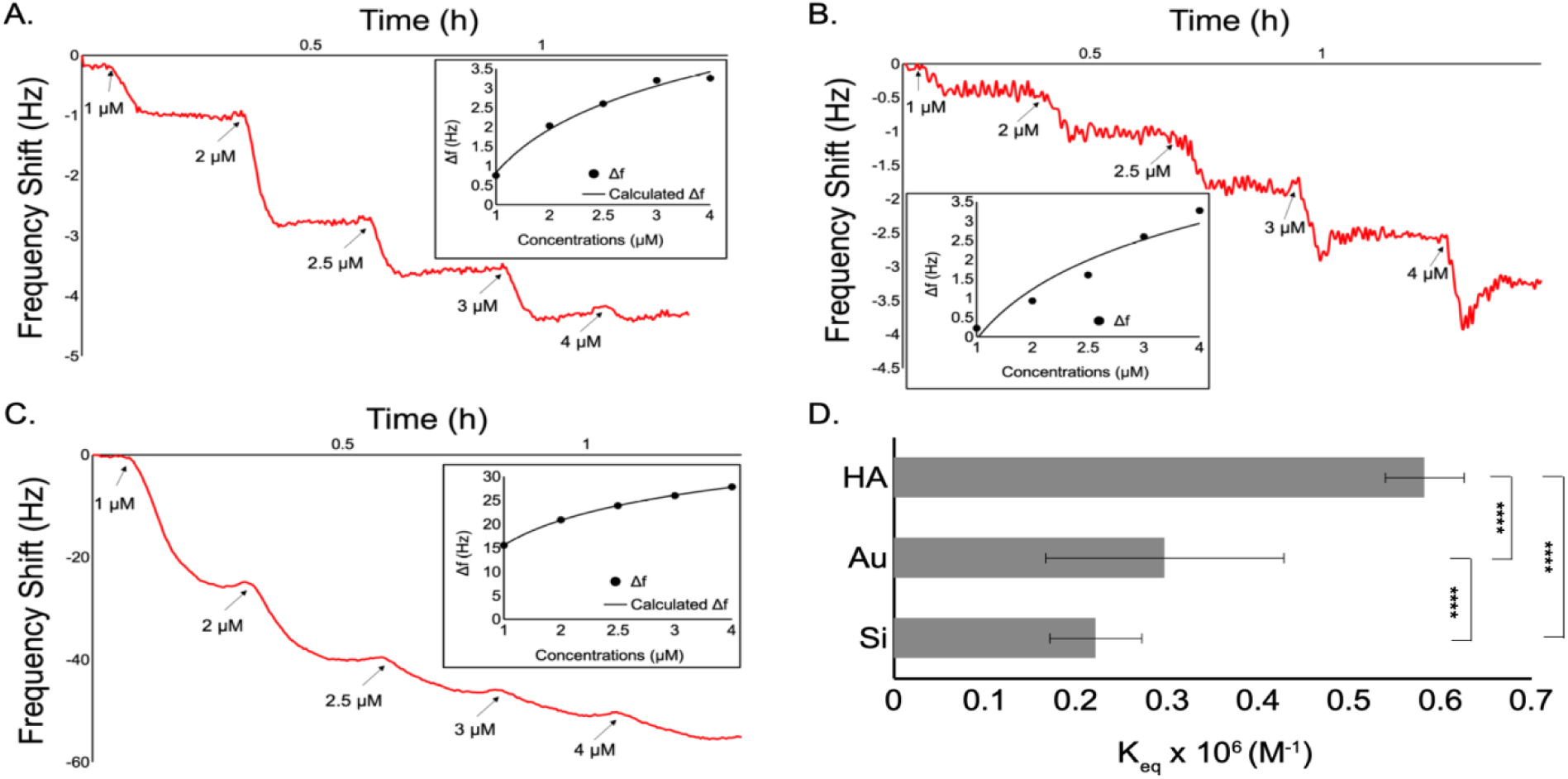
QCM measurement of aged CsgA-CsgB mix. Binding kinetics to the (A) gold, (B) silica and (C) HA coated quartz surface are shown as a resonance frequency change. D. Binding equilibrium constants of polymerized CsgA-CsgB mix on gold, silica and HA coated surfaces.

**Figure 5.**
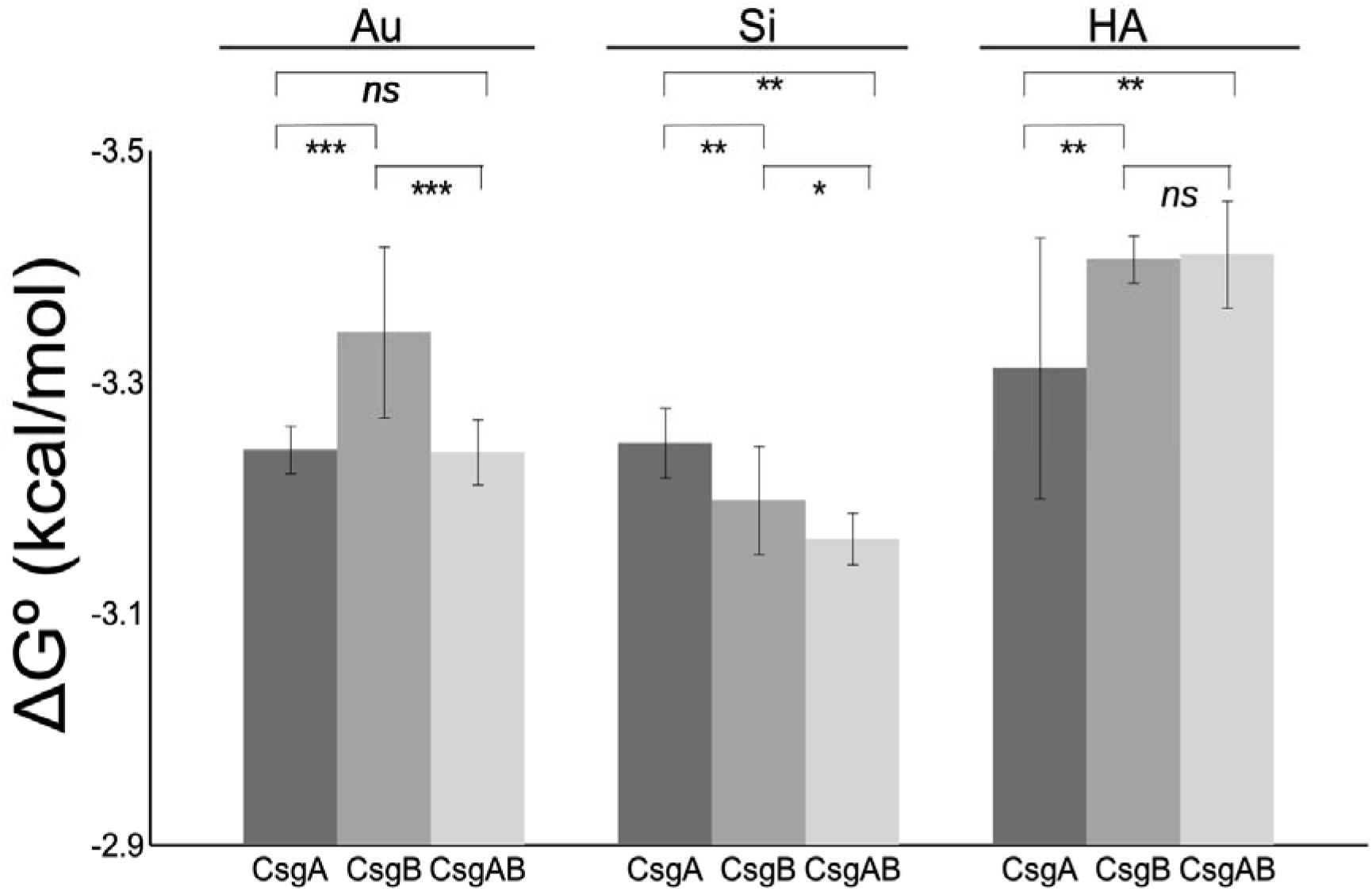
Binding free energy of polymerized CsgA, CsgB and CsgA-CsgB mix on gold, silica and HA.

In assessing surface binding characteristics, it should be noted that proteins and peptides vary in length, amino acid contents, as well as their charge (neutral or positive) and polarity or hydrophobicity. Based on diversity of peptides and proteins, it has been possible to identify common adsorption motifs on specific surfaces ^36^.

Another contributing factor in binding of a given protein is the properties of targeted surfaces. In a previous study, which examined the adsorption of CsgA subunit to graphene and silica surfaces by atomic simulation, it was reported that structural and sequence characteristics of CsgA affect the adhesive strength and consequently provide strong adhesion to both polar and non-polar surfaces ^24^.

Silica surface is composed of silanol groups (Si-OH) and siloxane bridges (Si-O-Si), and at pH values higher than 3 silanol groups tend to be deprotonated ^37^. Therefore, in our experimental setup, silica chips have a negative surface charge at pH 7. Surface charge is an essential parameter in terms of protein adsorption onto surfaces. The consensual view suggests that the reversible first step of protein adhesion onto silica results from the electrostatic properties of proteins ^38^. Therefore, at neutral pH, positively charged amino acids Arg, Lys, and His are considered essential for the electrostatic interactions with the negatively charged silica surface Interestingly, instead of the net charge of the protein, the total number of negatively and positively charged amino acids affect the binding affinity of proteins onto silica, probably due to increased polarity and hydrophilicity ^37^. Comparing our constructs in this study, CsgB contains more charged residues than CsgA. Even though we expect CsgB to have a higher affinity than CsgA, Figure 5 suggests the opposite. Here, considering the second step of protein adhesion can provide more detailed insight for those observations.

The second step, generally considered irreversible, depends on the ability of proteins to form structural changes on the surface. Tightly structured (or rigid) proteins do not induce conformational changes on the surface, so they are not very prone to adsorption. Proteins with weak internal interactions (or soft proteins), on the other hand, can structurally rearrange themselves on the surface more easily efficiently, and maximize the number of protein-surface interactions ^39^. In addition to the aforementioned electrostatic interactions, hydrophobic residues exposed to protein surface during the conformational changes may interact with the hydrophobic siloxane bridges on the silica surface. Therefore, in addition to the interaction with the charged residues, rigidity/softness of the protein structure is crucial for adhesion onto the silica surface. It has been demonstrated that curli fibers composed of only CsgB proteins exhibit more stiffness than curli fibers composed of only CsgA protein ^18^. Besides, combined secretion of CsgA and CsgB proteins in biofilms can produce mechanically stiffer nanofibers. Therefore, the stiffer nature of CsgAB fibers is in correlation with the behavior of mixture protein on the silica surface, which exhibits the highest Gibbs free energy, and therefore lowest silica affinity (Figure 5). Similarly, CsgA possessing the highest affinity to silica surface is in correlation with its lower stiffness compared to CsgB and mixture fibers. Therefore, the rigidity of the fibers appears to be a crucial parameter for adhesion onto the silica surface.

Hydroxyapatite (HA) is a naturally occurring mineral with the chemical formula Ca_5_(PO4)_3_(OH)]. HA has a high affinity for proteins and other biomolecules ^40^. Studies suggest that the pH and mechanical properties of proteins, as well as the surface topology of HA, can affect the protein-HA interactions ^41^. Computational studies suggested that –COO^−^ and -NH2 are the main interactions sites with HA and play crucial roles in the protein adsorption onto the HA surfaces by forming O-Ca-O connections and hydrogen bonds (H-bonds). Adsorption of proteins onto the Ca-rich HA surface can be mainly regulated by strong electrostatic interactions involving Ca atoms of HA and O atoms of carboxyl groups ^42^. Those strong electrostatic interactions also occur between PO_4_^-3^ anions in HA structure and the -NH^+^ and -C(NH)^+^ groups in the protein structure ^43^. Therefore, acidic amino acid residues Asp and Glu can serve as primary interaction sites with Ca, whereas basic amino acid residues Arg, Lys, and His can serve as interaction sites with PO4^-3^ anions. Overall, CsgA is more acidic (pI value 5.20) than CsgB (pI value 6.57), and CsgA has more acidic residues than CsgB. In contrast, CsgB has more basic amino acids than CsgA. ΔG calculations suggest that the interaction of CsgB with the HA surface is significantly higher than of CsgA. The strength of those interactions is ruled by interaction distances, as well as the number of interaction sites ^41, 42^. The higher affinity of CsgB can be due to the stronger interactions of basic amino acids with HA surface, rather than the number of interacting residues. The binding affinity of CsgB and CsgAB is not significantly different, indicating the forces that dominate CsgB interaction with HA is also governing CsgAB interaction with the surface.

Cysteines in proteins can interact with gold surfaces via irreversible chemisorption, however molecular dynamic simulations suggest that initial steps of recognition are due to electrostatic interactions. Computational studies suggested that histidine (His) and methionine (Met) residues have high gold binding propensity and can facilitate the initial adsorption of proteins onto gold surfaces, creating interactions approaching covalent bonds ^44^ Other molecular dynamics simulation studies proposed that aromatic residues form strong interactions with gold, possibly due to π electron mediated effects. Besides, positively charged amino acids also have strong binding affinities, whereas apolar aliphatic amino acids and negatively charged amino acids have weak interactions with the gold surface ^45^. Experimental studies showed that arginine (Arg), tryptophan (Trp), tyrosine (Tyr), and cysteine (Cys) amino acids are overrepresented in peptide groups with strong gold binding affinities ^46^.

CsgA **and** CsgB do not have any Cys in their structure; therefore, chemisorption of fibers onto gold surfaces is not affecting the binding affinity. Additionally, the number of aromatic groups in both CsgA and CsgB is the same; hence the aromatic group-Au surface interactions are not expected to contribute significantly to the difference in their surface binding affinity. To determine the charge of the proteins at pH 7.0, isoelectronic points were calculated using ExPASy. CsgA has a theoretical pI value of 5.20, and CsgB has a theoretical pI value of 6.57 that indicates both proteins are negatively charged at pH 7.0. CsgB has more positively charged amino acid residues than CsgA. Correspondingly, CsgB is expected to have a higher Au binding affinity than CsgA, because of the strong binding affinity of positively charged amino acids to Au. Besides, as the Met is reported to have a high gold binding propensity, five Met groups in CsgB can contribute to the Au binding affinity more than in CsgA, which has only two met groups in its structure. As anticipated, calculated ΔG values indicated that the affinity of CsgB to Au surface is significantly higher than of CsgA and CsgAB equimolar mix (p< .001). ΔG values of CsgA and CsgAB mix do not show a significant difference, indicating that the binding pattern of CsgA dominates the interaction of the mixture with the gold surface.

## Conclusions

CsgA and CsgB proteins, which are excellent candidates for nanomaterial synthesis due to their self-assembly capabilities, and can be easily manipulated by genetic engineering.

Depending on the intended purpose, binding of proteins to selected surfaces at controlled and adjustable levels is essential in the design and development of various amyloid-based functional materials. To obtain controlled and adjustable binding, it is necessary to examine and characterize systems, including proteins and target surfaces.

Here we reported that CsgA, CsgB and their mixture behave differently in terms of binding and interaction with gold, silica and HA surfaces. According to our observation CsgB protein is the key protein that dictates the final adhesiveness of the final CsgAB protein mixture on HA and silica surfaces. In contrast to this observation CsgB, however, is not the key component in controlling the behavior of CsgAB mixture on gold surface. Our findings provide an understanding of binding behavior of CsgA, CsgB and their mixture to gain a better control on their adhesion on a targeted surface.

## ASSOCIATED CONTENT

### Supporting Information

Details of cloning experiments, and protein purification

## ACKNOWLEDGMENT

We thank TUBITAK for the financial support through Grant No: 216M127, and TÜBA-GEBIP.

